# The enigma of the SARS-CoV-2 microcirculation dysfunction: evidence for modified endothelial junctions

**DOI:** 10.1101/2023.04.24.538100

**Authors:** L Bouillet, M Benmarce, C Guérin, L Bouvet, O Garnier, D K Martin, I Vilgrain

**Affiliations:** Univ. Grenoble Alpes, CNRS, TIMC-IMAG / T-RAIG (UMR 5525), 38000 Grenoble, France; Univ. Grenoble Alpes, INSERM U13, CEA, Institute of Interdisciplinary Research of Grenoble (IRIG), Laboratory of Biosciences et Bioingénierie pour la santé (BGE)-Biomics, 38000 Grenoble, France; Grenoble Hospital Grenoble Alpes (CHUGA), Univ. Grenoble Alpes, 38000 Grenoble, France; Univ. Grenoble Alpes, CNRS, TIMC-IMAG / SyNaBi (UMR 5525), 38000 Grenoble, France

**Keywords:** Endothelium, Cadherin, Pulmonary, Post-translational, Physiopathology, Metallopeptidase

## Abstract

Published evidence indicates that Severe Acute Respiratory Syndrome-Corona Virus (SARS-CoV-2) infection causes endothelial cell (EC) injury in the Coronavirus Disease 2019 (COVID-19). Endothelial junctions (EJ) are crucial to maintain EC integrity and normal microvascular functions due to the adhesive properties of Vascular endothelial (VE)-cadherin to glue EC together. Here we report studies in vitro and in vivo that indicate VE-cadherin to be a target for cleavage by ACE2. We have identified that the extracellular domain of VE-cadherin contains these two amino acid sequences at the positions ^256^P-F^257^ and ^321^PMKP-^325^L for ACE2 substrate recognition. Incubation of purified sVE with ACE2 revealed a dose-dependent loss of immunoreactivity detected with an antibody directed against the Extracellular domain 1 (EC1) domain of sVE. We confirmed the presence of ACE2 on ECs using immunofluorescence studies, and by western blotting on ECs extracts. We also present evidence from patients with severe COVID-19 disease for a circulating form of ACE2. Its apparent molecular weight of 70 kDa is in agreement with a previously described extracellular form of ACE2 bearing the catalytic site of the ectopeptidase. Consistent with the experimental evidence for our hypothesis, the level of circulating soluble VE-cadherin fragments was increased in the blood of patients with severe COVID-19 disease. Further studies are needed to determine if increased circulating fragments of ACE2 and VE-cadherin may contribute to the future development of post-acute COVID-19 syndrome characterized by vascular endothelial injury, hypoxia, and inflammatory state.

**Impact Statement:** SARS-CoV-2 infection promotes vascular dysfunction but the processes are not completely understood. The vascular endothelium is composed of a monolayer of endothelial cells (ECs) that exclusively express VE-cadherin at adherens junctions (AJs). The published structure of VE-cadherin has revealed crucial residues in the domains EC1-2 for ECs adhesiveness. In this report, we demonstrate for the first time that VE-cadherin is a target for ACE2 ectoenzyme in the domains EC2-3. In addition, in COVID-19 patients’ blood, we identify truncated forms of ACE2 and VE-cadherin that are correlated with severe SARS-CoV-2 infection. Because the turnover rate of ECs is very low, this could provide part of the explanation for Long CoVID-19 disease. These exciting results highlight the role of proteases and AJs, and the need for continuing efforts to elucidate whether these circulating proteins might be of prime significance for clinicians to facilitate personalized medicine.

## Introduction

The world continues to deal with successive waves of the Coronavirus disease 2019 (COVID-19), for which often disabling sequelae are called “post-COVID-19 syndrome”. It has a worldwide impact and was first diagnosed in 2019, in Wuhan, China^1^. Up to now, more than 280 million cases have been confirmed, leading to over 5.4 million deaths in the world^2^. Among the distinctive features of COVID-19 are the associated vascular changes, which might be due to direct endothelial infection. Evidence for endothelial infection was reported by Varga et al^3^ who found viral bodies in endothelial cells (ECs) from glomerular capillary loops of the transplanted kidney of a patient, and endothelialitis in an additional two patients. Varga et al^3^ concluded that all three patients had an endothelial cell infection, diffuse endothelial inflammation, and thus endothelium destabilization. Ackermann et al^4^ recently observed morphologic and molecular changes in the peripheral lung of patients who died from COVID-19, which consisted of severe endothelial injury, widespread thrombosis with microangiopathy, and vascular angiogenesis. Chhetri et al^5^ showed that many COVID-19 patients exhibited severe metabolic acidosis, which indicated possible microcirculation dysfunction. Therefore, understanding the mechanism of endothelial dysfunction in COVID-19 is warranted for exploring better clinical care for patients.

Our hypothesis regarding SARS-CoV-2 infection in COVID-19 disease considers the physiology of the vascular endothelium. The integrity of the dense vascular network surrounding small pulmonary alveoli depends upon endothelial adherens junctions and the pericyte coverage of the small capillaries. Endothelial junctions insure the tight association between ECs because of the strong adhesive properties of a cell adhesion transmembrane protein called Vascular Endothelial (VE)-cadherin exclusively expressed in ECs^6^.The whole VE-cadherin structure maintains EC adhesion by homotypic interactions between the five homologous repeats each of about 110 residues that are stabilized by binding of Ca^2+^ ions^6^. The cytoplasmic domain of the protein is bound with the catenins that form the structural bridges with the actin cytoskeleton to enhance adhesion. The pivotal role of VE-cadherin in vascular physiology has been demonstrated in deficient embryos who died at 10.5d from extensive angiogenesis defects^7)^. Because of these VE-cadherin extracellular and cytoplasmic interactions, the endothelial junctions are extremely stable in adults except upon challenge by inflammatory mediators, such as histamine, thrombin, VEGF, TNF-α, and platelet-activating factor. Following such challenge, the covalent modification of VE-cadherin by tyrosine phosphorylation is associated with the dissociation of junctions and increased EC permeability^8-13^. Indeed, the covalent modification of VE-cadherin precedes the cleavage of its extracellular domain to release soluble VE-cadherin (sVE) into human blood. A clinical example is in Bradykinin-mediated Angioedema patients (HAE) during an attack, where Bradykinin is the major mediator of the vascular leakage that causes recurrent swellings, and sVE is found in the blood^13,14^. At both cellular and molecular levels, we demonstrated that the release of sVE was the result of Bradykinin-induced VE-cadherin phosphorylation as well as due to the direct proteolytic action of Kallikrein. Indeed, this protease has two consensus sites of cleavage at the peptide bonds ^411^R-T^412^ and ^565^R-T^566^ in the VE-cadherin extracellular adhesive domain (sequence identifier, P03952). In newly diagnosed Rheumatoid Arthritis (RA) patients (RA), the blood sVE level resulting from the action of TNF-α on ECs was correlated significantly with the Disease Activity Score at baseline and after 1-year follow-up^8^. In addition to RA and HAE, patients with systemic inflammation and sepsis also had sVE in the blood^15^. Altogether, these data collectively support that cytokine-induced structural modifications of VE-cadherin lead to endothelial instability and capillary leakage. Clinically significant support for this hypothesis was reported by Mehta et al^15-17^, who described a cytokine storm as a result of the SARS-CoV-2 infection, an increased capillary leakage of fluid, and recruitment of immune-inflammatory cells in lungs. Because of the large number of proinflammatory cytokines in the COVID-19 environment, we suggest that all of these can target the endothelial junctions.

It is important to note that SARS-CoV-2 enters human cells predominately by binding to the angiotensin-converting enzyme 2 (ACE2)^18,19^. In vivo, ACE2 is expressed in ECs, renal tubular epithelium, vascular smooth muscle cells, and pericytes^20,21^. For example, alveolar pneumocytes in the lung are a possible site of entrance for SARS-CoV-2 since type I and II pneumocytes express ACE2. Of great significance is that ACE2 is a metallopeptidase orientated with its catalytic site facing the extracellular space as an ectoenzyme, and thus it can metabolize extracellular proteins. Its site of cleavage has been mapped and corresponds to the amino acid sequence PMKP–L, where the bond P-L is cleaved (24). Active soluble ACE2 has been detected in the plasma of patients with heart failure^22,23,25^, which suggests that its diffusion throughout the body via the bloodstream might also contribute to abnormalities of microcapillaries leading to endothelial inflammation, and thus endothelium destabilization^22^. Altogether, these data from the literature and our previously published data strongly suggest that VE-cadherin, the adhesive protein in charge of vascular integrity, might be the target of proteases activated in the COVID-19 microenvironment involved in vascular dysfunction.

The aims of the present study were to determine whether the extracellular domain of VE-cadherin was a substrate for ACE2 in vitro, to examine the potential expression of ACE2 in ECs, to analyze if there were circulating forms of ACE2 in blood from COVID-19 patients with mild or severe disease, and to analyze if soluble VE-cadherin was released into the blood of those COVID-19 patients.

## Materials and Methods

### Antibodies

Recombinant Human ACE2 Protein (933-ZN) was purchased from R&D Systems (Minneapolis, MN, U.S.A.) and stored in a buffer according to the manufacturer’s instructions. Monoclonal mouse anti-human ACE2 Antibody (MAB933) was purchased from R&D Systems (Minneapolis, MN, U.S.A.). VE-cadherin extracellular domain (EC1-5) was produced in the laboratory. Peroxidase AffiniPure Goat Anti-Mouse IgG (H+L) (AB-2338447), Peroxidase AffiniPure Goat Anti-Rabbit IgG (H+L) (AB-2307391), and Cy3™ AffiniPure Goat Anti-Mouse IgG (H+L) (AB-2338680) were all purchased from Jackson Immunoresearch (Ely, Cambridgeshire, U.K.). Hoechst solution (33258) was purchased from Sigma Aldrich. Mouse monoclonal BV9 anti-human VE-cadherin antibody and rabbit polyclonal anti-EC1 anti-human VE-cadherin antibody were produced in the laboratory.

### Cell culture

Human Umbilical Vein Endothelial Cells (HUVECs) were grown onto fibronectin-coated plates to reach confluency or 4 to 7 days in M199 medium supplemented with 10% fetal calf serum (FCS) and 2% low serum growth supplement (Cascade Biologics, Portland, Oregon, U.S.A.). Experiments were performed using early cell passages, from P0 to P5. Human embryonic kidney cells (HEK293-EBTNA were grown as described in^12^.

### Immunofluorescence

HUVECs were grown on plates until confluence then fixed with 3.5% (w/v) paraformaldehyde in PBS. Permeabilization or not was performed with methanol at -20°C for 10 min followed by incubation in Triton X-100 (0.1% in PBS) for 10 min. After three washes with PBS, the cells were then incubated for 1 h with the rhodamine-labeled secondary antibody. After three washes in PBS, the nuclei were labeled with 1 μg/ml Hoechst for 5 min and then washed three times with PBS. The coverslips were then mounted with Aquamount mountant. Fluorescence images were collected using ApoTome Fluorescence Microscope.

### Proteins extraction, SDS-PAGE and Immunoblotting

HUVECs were harvested and then suspended with lysis buffer (NaCl 150mM, Tris 20mM, HCl pH 7.4, Sucrose 270mM, EDTA 1mM, EGTA 1mM, Na3VO4 2mM, NaF 5mM, DTT 1mM, Triton 0.5%). Cell lysates were homogenized, sonicated for 5 seconds and centrifuged at 14,000 rpm for 10 min the supernatants were stored at -80°C until used. Total extracted proteins were identified using the micro-bicinchoninic acid method. Protein extracts were stored at -20°C before use.

The human glycosylated VE-cadherin extracellular domain was produced from human embryonic kidney cells (HEK293-EBTNA) according to a previously published protocol^26^.

Proteins were subjected to electrophoresis on SDS-PAGE 10%, and then transferred onto a 0.45μm nitrocellulose membrane. Membranes were blocked with milk, incubated with primary antibodies and corresponding HRP-conjugated secondary antibodies. Immunoreactive proteins were visualized by Bio-Rad ChemiDoc™ with ECL reagent (Bio-Rad Laboratories) and were quantified using densitometry with Image-J software (NIH, Bethesda, MD). All Western blots are representative of at least three independent experiments with similar findings.

### Patient Sera

A serum bank was available at Grenoble University Hospital. Analyses were performed on samples from 9 patients with COVID-19. Five of them had a severe form of COVID-19 with a passage in resuscitation and intubation. The other 4 patients had mild forms, requiring only oxygen therapy.

### Statistical Analysis

All the experiments were repeated at least three times. Values represent the mean ± standard deviation of three determinations from three different wells or dishes in the same experiment. Data are expressed in arbitrary units as the mean ± SD of at least 3 identical experiments and were compared using Mann-Whitney test. For all tests, *P* values less than or equal to 0.05 were considered significant.

## Results

### Endothelial cell junctions and VE-cadherin

A small capillary is typically composed of three endothelial cells (EC) surrounded by pericytes (P) (**Fig 1A**). The integrity of the vascular network depends upon endothelial adherens junctions (EJ). The overlap of two ECs is observed by transmission electron micrograph and represents the adherens junction (**Fig 1B)**. The complete sequence of VE-cadherin is presented in **Fig 1C** (sp P33151). The extracellular domain is highlighted in yellow. The amino acids in red represent the consensus sequences recognized by the ACE2 ectoenzyme. The 5-domain structure of the extracellular fragment of VE-cadherin extracellular domain is shown in **Fig 1D** with the localization of the epitopes of the antibodies EC1 and BV9 directed against human VE-cadherin. We purified recombinant protein from conditioned media from human embryonic kidney cells. The coomassie-stained SDS-PAGE gel showed that recombinant extracellular domain of VE-cadherin exhibits an apparent molecular weight of 90 kDa (**Fig 1E, left panel**). This is the apparent molecular weight of the protein since is normally glycosylated. The glycosylation was confirmed by a treatment of the protein with PGNase, which resulted in an apparent molecular weight of 75kDa (**Fig 1E, right panel**). The monoclonal mouse anti-human VE-cadherin recognized the recombinant protein in a dose-dependent manner, which further confirmed that the recombinant protein was the human glycosylated extracellular domain of VE-cadherin (**Fig 1F**). The precise molecular mass of the purified protein was determined by MALDI-TOF mass spectrometry to be 430 μg/mL. Altogether, this allowed us to study whether VE-cadherin extracellular domain was a substrate for ACE2 in vitro.

**Figure 1:**
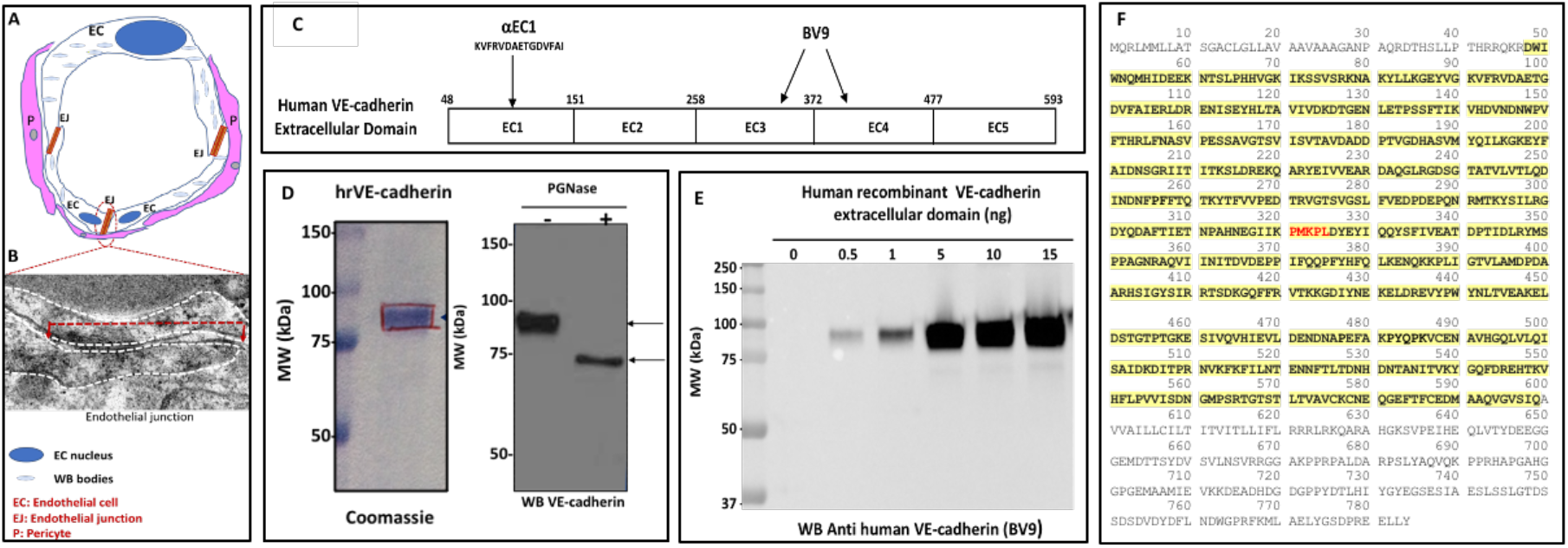
Endothelial junction and sVE-cadherin. **A**. Schematic representation of a capillary composed of three endothelial cells (EC). The capillary is surrounded by pericytes (P). The overlap of two ECs is the adherens junction (EJ) **B**. Electron microscopy of the endothelial junction. The dashed white line delineates the contour of two ECs. **C**. Amino acid sequence of human VE-cadherin. The yellow color highlights the amino acid sequence of the extracellular domain of the protein. The amino acid sequence of the ACE2 putative cleavage site is highlighted in red. **D**. Schematic representation of VE-cadherin extracellular domain with the localization of the epitope of the mouse monoclonal antibody directed against human VE-cadherin. **E**. Coomassie-stained SDS-PAGE gel showing that human recombinant extracellular domain of VE-cadherin exhibits an apparent molecular of 90 kDa and immunoblotting of VE-cadherin not treated (left lane) or treated (right lane) with PGNase. **F**. Immuno blotting of human recombinant VE-cadherin antibody (BV9).

### VE-cadherin extracellular domain is a substrate for ACE2

Human ACE2 (angiotensin-converting enzyme 2) is a transmembrane zinc-containing metalloenzyme of 805 amino acid residues, with seven N-glycosylation sites and three disulfide bridges (Swiss-Prot code Q9BYF1). It is a single-pass type I integral membrane glycoprotein, orientated with the N-terminus and the catalytic site facing the extracellular space (an ectoenzyme), where it can metabolize circulating peptides. Since VE-cadherin is expressed as a transmembrane protein in ECs, its large extracellular domain could represent a potential substrate for ACE2 ectoenzyme since VE-cadherin presents post-translational modifications under cytokine challenge. We tested this hypothesis using a commercially available human recombinant ACE2 enzyme and the human recombinant extracellular domain of VE-cadherin. First of all, the native ACE2 ectoenzyme was analyzed by SDS-PAGE and immuno blotting using the antibody directed against the ectodomain of the enzyme. The ACE2 ectoenzyme was detected in the range from 5 to 10 ng and exhibited an apparent molecular weight of 110 kDa (**Fig 2A**). Then, the enzymatic reaction with the extracellular domain of VE-cadherin was tested in vitro using the prediluted ACE2 in 50 mM 2-(N-Morpholino)-ethane sulfonic acid (MES), pH 6.5, 300 mM NaCl, 10 µM ZnCl2, 0.01% Brij 35. Increasing concentrations of ACE2 (0, 0.3, 3, 10, 30, 100, 300, 1000 ng) were added to the extracellular domain of VE-cadherin (40 ng) overnight at room temperature. The mixture was then analyzed by SDS-PAGE and western blotting with the rabbit polyclonal EC1 antibody directed against the EC1 domain of the extracellular domain of VE-cadherin (**Fig 2B**). As a result, the immunoreactivity of the 90 kDa band corresponding to the VE-cadherin extracellular domain decreased in a dose-dependent manner after ACE2 incubation. Densitometric analysis of the 90 kDa band showed a dose-dependent decrease in VE-cadherin band density starting at 10 ng of ACE2 to decrease progressively between 100 and 300 ng (**Fig 2C**). Altogether, this suggests that the extracellular domain of VE-cadherin is a target for ACE2 ectoenzyme.

**Figure 2:**
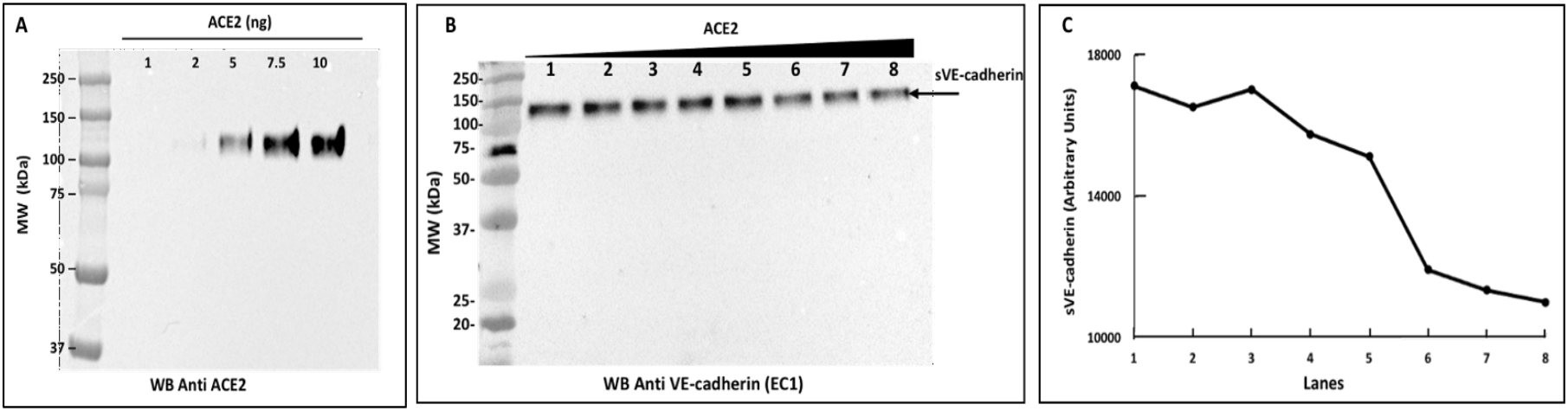
The extracellular domain of VE-cadherin is a substrate for the protease ACE2 : **A**. Commercially available human recombinant ACE2 (1 to 10 ng) was analyzed by immunoblot with the primary monoclonal anti-human ACE2 antibody followed by a secondary antibody anti-mouse IgG (1:2500) overnight. **B**. VE-cadherin extracellular domain (40 ng) was incubated overnight at room temperature with increasing concentrations of ACE2 (0, 0.3, 3, 10, 30, 100, 300, 1000 ng-Lane 1 to 8) in 50mM 2-(N-Morpholino)-ethane sulfonic acid (MES), 300 mM NaCl, 10 μM ZnCL2, 0.01 % Brig 35, pH 6.5. The reaction was stopped with Laemmli buffer.The samples were analyzed by SDS-PAGE and immunoblotting. The soluble VE-cadherin was then detected with the rabbit polyclonal anti-EC1 antibody (1:250) directed against the EC1 domain of human VE-cadherin and by the secondary antibody anti-rabbit IgG (1:1000). **C**. Densitometric analysis of the 90 kDa using ImageJ software quantification showed a dose-dependent decrease in VE-cadherin band density. This experiment is representative of three independent experiments.

### Detection of ACE2 in ECs

In an attempt to determine the ACE2 expression on ECs, we performed immunofluorescence studies on HUVECs as a proof of concept because these are primary cultures that do de-differentiate and lose the expression of proteins over several passages. Indeed, VE-cadherin was clearly observed in these cells with a monoclonal antibody that recognized the extracellular domain of the protein of VE-cadherin (BV9) (**Fig 3A)**. Using then the antibody directed against the ectdomain of the ACE2, we clearly identified that ACE2 ectoenzyme was expressed at the surface of the HUVECs **(Fig 3B)**. Note that the immunofluorescence protocol was used without a permeabilization step, which further supported the interpretation that ACE2 is expressed on the surface of the cells.

**Figure 3:**
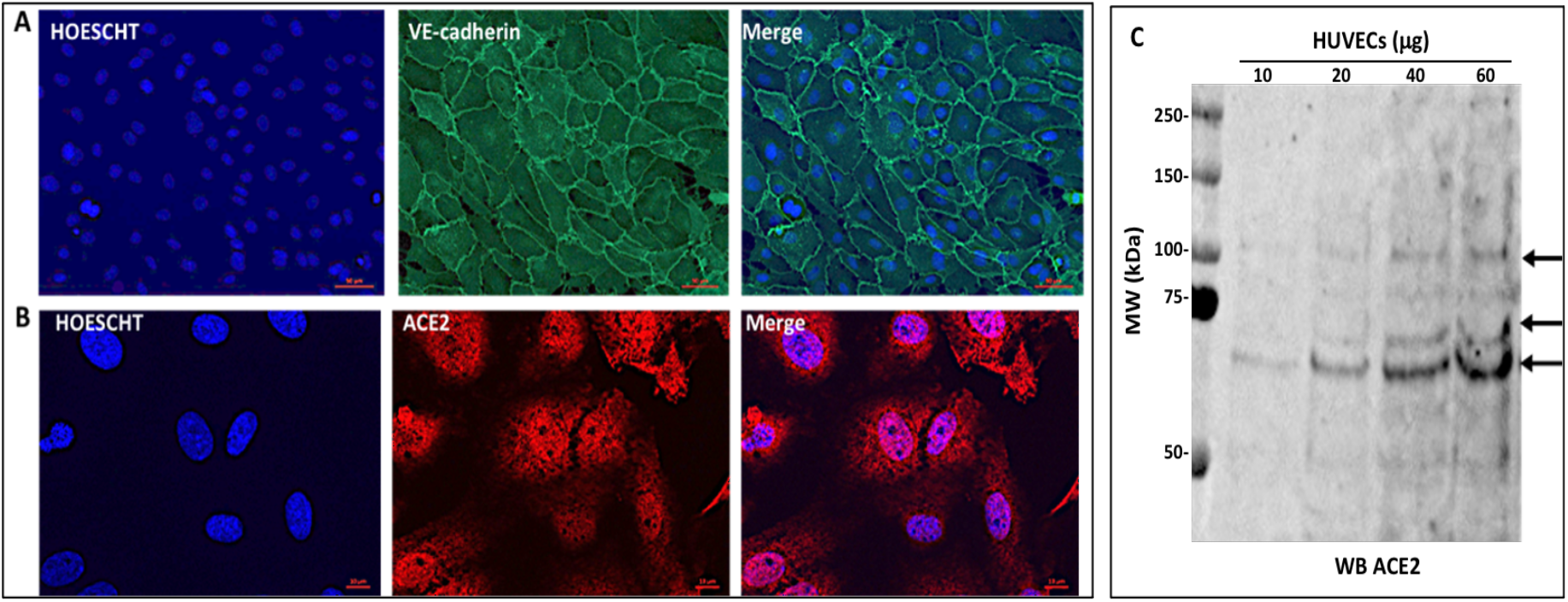
Immunodetection of VE-cadherin and ACE2 in HUVECs. **A :** Immunofluorescence of VE-cadherin with BV9 antibody (2 μg/mL). (Scale bar 50μm). **B** : Immunofluorescence of ACE2 (red) using an antibody that recognize the ectodomain of the protein (anti-ACE2 10 μg/mL) (Scale bar 10μm). **C**. ACE2 detection in HUVECs lysates (10, 20, 40, 60 μg) by immunoblotting with the anti-ACE2 antibody (10µg/mL). Fluorescence images were taken with x40 objective. This experiment is representative of four independent experiments.

We next examined the ACE2 expression at the protein level in HUVECs. The cells were harvested in a lysis buffer that contained 0.5% Triton-X100, and were then sonicated for 30 sec on ice to extract the membrane proteins. The homogenates were centrifuged at 14,000 rpm for 10 min. The supernatant was assayed for protein content and increasing quantities of the extracted proteins were analyzed onto a 10% SDS-PAGE gel and immunoblotting with the same ACE2 antibody that was used for immunofluorescence experiments. Results showed that HUVECs expressed three bands with an apparent molecular weight around 110 kDa, 70 kDa, and 60 kDa respectively (**Fig 3C**). The 110 kDa fragment corresponded to ACE2 since it has the same molecular weight as the control (cf. **Fig 2A**) and the other fragments most likely are the result from the cleavage of ACE2 by proteases in HUVECs (27).

### Detection of circulating ACE2 in blood samples from COVID-19 patients

A disintegrin and metallopeptidase domain 17 (ADAM17)/tumor necrosis factor α-converting enzyme cleaves ACE2 from the cell surface^(27)^. This produces circulating ACE2, an ectoenzyme with an active ectodomain. We next wanted to determine if patients with a mild or severe COVID-19 disease have the circulating form of ACE2 in the blood. The serum obtained from the blood samples of these COVID-19 patients was diluted serially according to our published protocol^12^. The diluted sera were then analyzed on a 10% SDS-PAGE and immunoblotting with the anti-human ACE2 ectodomain monoclonal antibody. As shown in the **Fig 4A**, the presence of an ACE2 fragment was detected in the blood from patients with a severe SARS-CoV-2 infection while it was barely detectable in patients with mild disease (**Fig 4B**).

**Figure 4:**
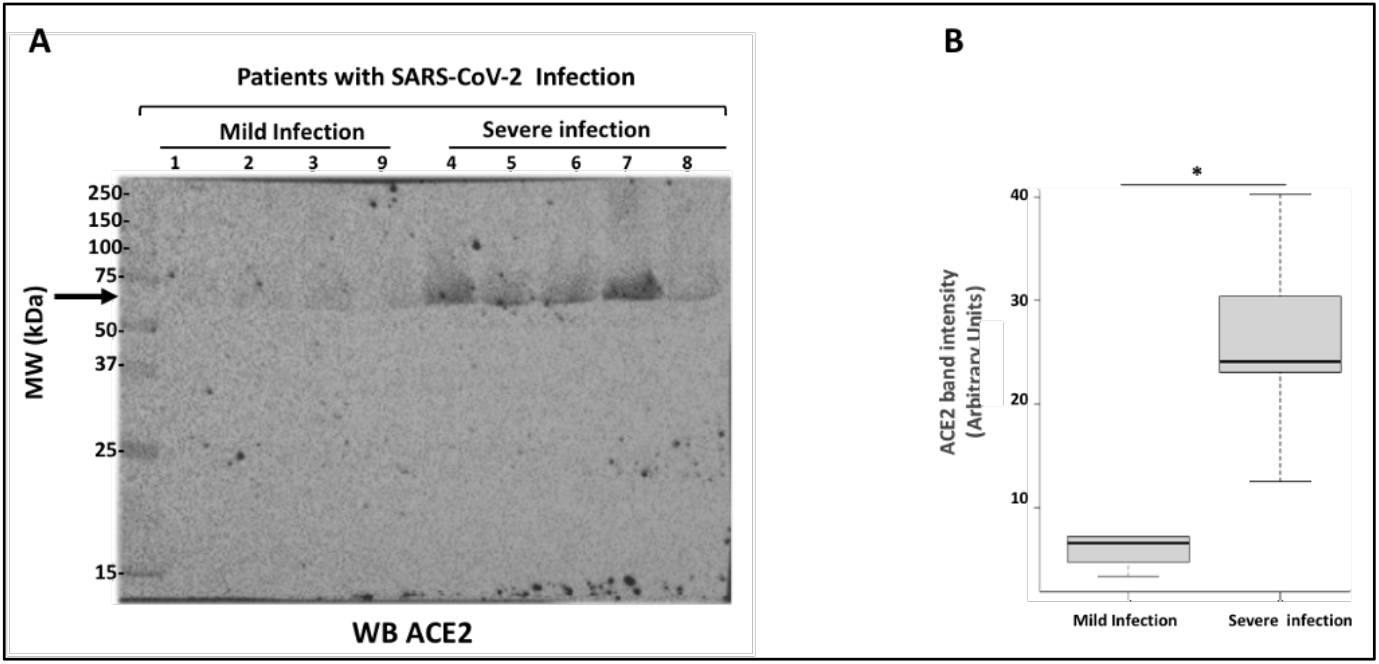
Analysis of circulating ACE2. **A :** Blood samples from COVID-19 patients were analyzed by SDS-PAGE and immunoblotting. Blood samples were diluted serially from 1 to 10 followed by a dilution of 1 to 2.5. 15μL of sample preparations were loaded onto a SDS-PAGE and then transferred onto cellulose membrane. The membranes were then incubated with the anti-ACE2 antibody (5μg/mL) followed by incubation with the anti-mouse peroxidase antibody. All experiments were performed in triplicate. **B**. Densitometry values allowed to compare the groups (n=4 for mild infection and n=5 for severe infection). Error bars represent mean ± SE of means. P values from analysis of variance were assessed using the Mann-Whitney test (*p <0.05).

### Analysis of soluble VE-cadherin in blood samples from COVID-19 patients

Then we determined whether soluble forms of VE-cadherin were detected in the blood of the same patients using the antibody directed against the extracellular domain EC1. As shown in **Fig 5A**, several fragments of VE-cadherin were detected in patients with either the mild or severe forms of COVID-19. In the patients with a mild form of the disease, only a band with an apparent molecular weight of 54 kDa was detected. In patients with the severe SARS-cooV-2 infection, three bands were detected with an apparent molecular weights of 70, 62, and 54 kDa (**Fig 5B**). Taken together, these results show that depending on the severity of the disease the soluble VE-cadherin seems to be a good clinical biomarker.

**Figure 5.**
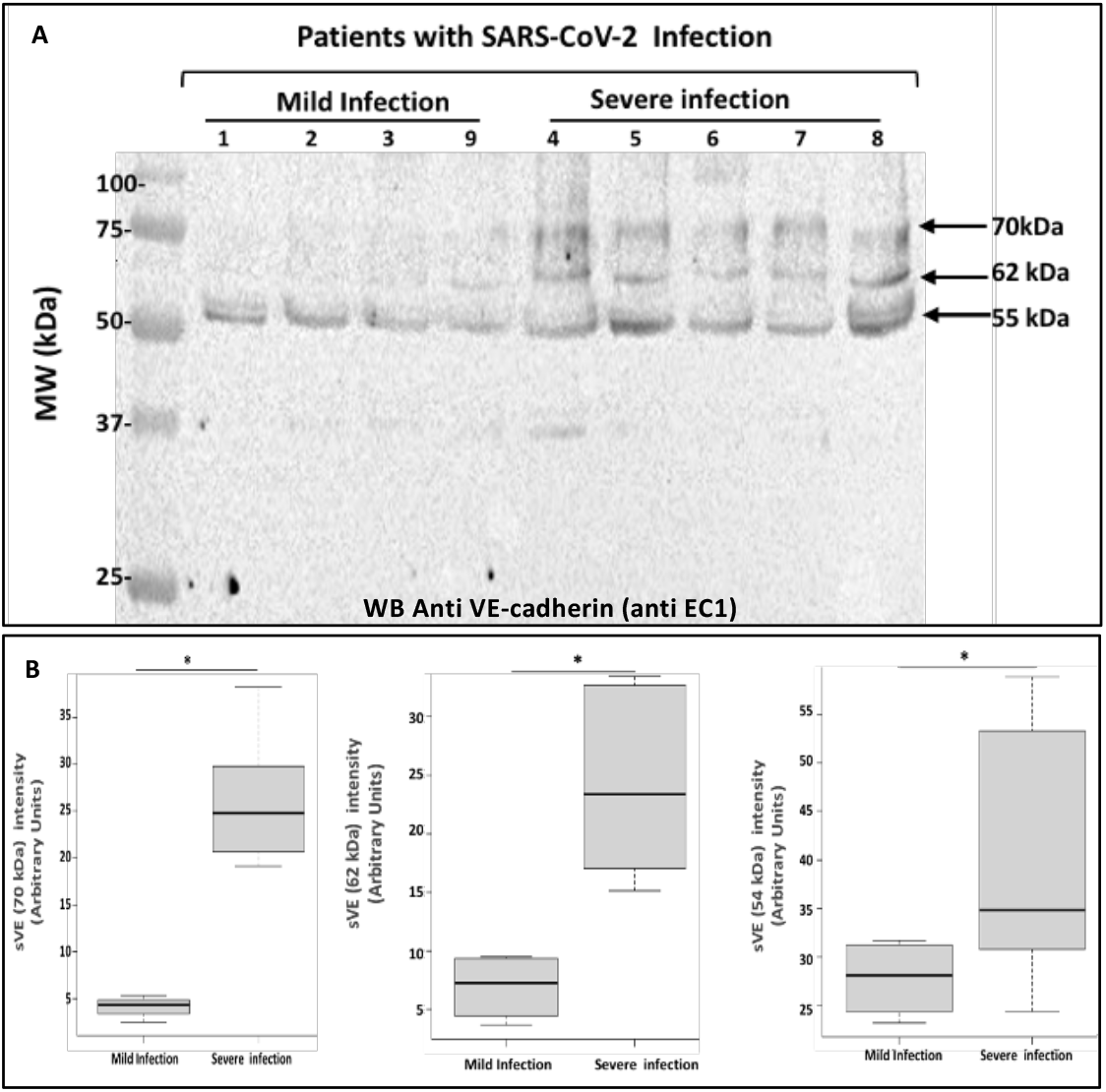
Analysis of sVE-cadherin in blood from COVID-19 patients. **A**. Blood samples from COVID-19 patients were analyzed by SDS-PAGE and immunoblotting. Blood samples were diluted serially from 1 to 10 followed by a dilution of 1 to 5 (total dilution 1/50). 15μL of sample preparations were loaded onto the gel and then immunoblotted with the anti-VE-cadherin antibody (EC1 1/250) overnight at 4°C followed by an incubation with the goat anti-rabbit peroxidase antibody (1:2000). This experiment is representative of three independent experiments. **B**. Densitometry values allowed to compare the groups (n=4 for mild infection and n=5 for severe infection). P values from analysis of variance were assessed using the Mann-Whitney test (*p <0.05).

### Proposed scheme model for endothelial aggression following SARS-CoV-2 infection

We have illustrated in **Fig 6** a scheme of the COVID-19 microenvironment at the pulmonary level. The very close interaction between pneumocytes (P1, P2), alveolar macrophages, and EC of pulmonary capillaries led us to propose that the infection by the virus creates an environment where several actors (cytokines, proteases) can induce structural changes in VE-cadherin (eg TNF) such as the phosphorylation processes and cleavage of its extracellular adhesive domain. Interestingly, it is noteworthy that SARS-CoV-2 enters human cells mainly by binding to ACE2. Thus we suggest that the protease ACE2 has a substrate just located in the membrane of ECs and can cleave VE-cadherin. Because active ACE2 has been recently detected in plasma, its diffusion in the body through bloodstream might also be part of abnormalities of microcapillary leading to endothelial inflammation, and thus endothelium destabilization.

**Figure 6:**
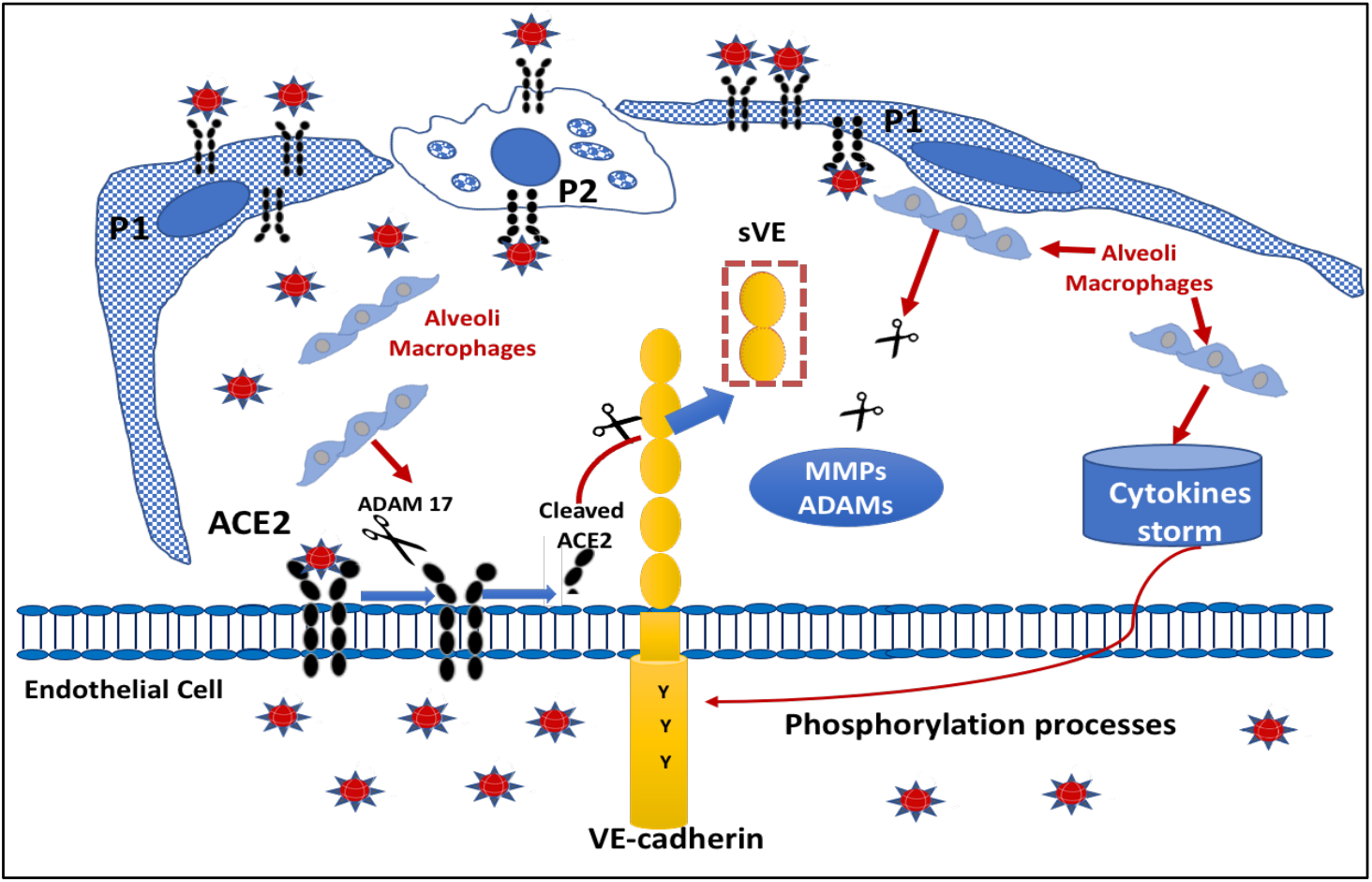
Schematic model of Pulmonary Endothelial aggression in COVID patients: **A**. In the pulmonary environment, the endothelial cells (EC) are surrounded by pneumocytes (P1, and P2) and macrophages that are responsible for the cytokine storm. ACE2 is present on EC as a transmembrane protein whose catalytic site is outside the cells (ectopeptidase). ACE2 can be cleaved by ADAM 17 and leads to the generation of a circulating active form of ACE2. VE-cadherin is a transmembrane protein exclusively expressed in ECs which can be subjected to post-translationnal modifications including tyrosine phosphorylation upon cytokine challenge. This covalent modification will lead to a conformational change in the protein structure which presents a high sensitivity to proteases. Thus the ACE2 enzyme will act directly to its potential site of cleavage generating the fragments of VE-cadherin seen in the patient’s blood.

## Discussion

The clinical reports coupled with previous publications provide strong evidence for EC infection and endothelialitis following SARS-CoV-2 infection. According to the Zurich team^4^, SARS-CoV-2 infection causes vascular inflammation in different organs. It is important to note that SARS-CoV-2 infects human cells mainly by binding to the protease called ACE2. Our central hypothesis is that ACE2 can lead to the cleavage of the extracellular domain of VE-cadherin at specific ACE2 target sequences, which then destabilizes the EJ and causes microcirculation dysfunction. This hypothesis has never been considered previously.

We produced the fully glycosylated extracellular domain of human VE-cadherin (sVE) to test whether it was an atypical substrate for ACE2. As a proof of concept, we incubated sVE overnight with ACE2 in the presence of Zn to catalyze the enzymatic reaction^24^. With the use of the anti-EC1 antibody, the detection of the immunoreactive band was decreased by approximately 35% compared to that detected with BV9. This suggests that, because the putative site for ACE2 is located in the middle of the EC3 domain, the fully glycosylated extracellular domain of human VE-cadherin (sVE) has lost some immunoreactivity and has been cleaved by ACE2. The EC1 epitope has been shown to form the homophilic binding of VE-cadherin and mediates homophilic adhesion in the vascular endothelium^27^. Classical cadherin adhesive interfaces have been localized to the EC1 domain, and the adhesive binding is strongly affected by the Ca^2+^ binding sites between the EC1 and EC2 domains. The adhesive binding of VE-cadherin is among the strongest of the type II cadherins. Nonetheless, it remains to be determined whether VE-cadherin employs an alternative *cis* interface between neighboring proteins. Domain shuffling experiments have shown that the determinants of cellular specificity reside in the EC1 domain and often display homophilic dimerization^27^. We acknowledge that our study has limitations. In our in vitro experiments of cleavage of VE-cadherin by ACE2, we did not detect a protein fragment that was susceptible to be detected by the anti-EC1 antibody. The two putative reasons for that are (i) the cleaved protein fragment is in a low amount and is below the limit of detection by the antibody, and (ii) the degradation product is in a conformation that does not allow the recognition by the antibody likely because of the glycosylation sites of the protein. Further investigations will be required to address those issues.

In concurrence with the literature^22^, we demonstrated by immunofluorescence and by immunoblotting that ACE2 is expressed on HUVECs. The expression of ACE2 was at the surface of the HUVECs, where it can function as the primary receptor for SARS-CoV-2. For example, it may facilitate fusion and endocytosis of SARS-CoV-2 into the pulmonary ECs through the interaction of the viral spike protein with the membrane-bound ACE2. We measured that the ACE2 protein extracted from the HUVECs (110, 70, 60 kDa), which suggests that it was cleaved during the process of extraction by other proteases in the ECs. A likely cause for such cleavage is ADAM17 (a disintegrin and metallopeptidase domain 17), which was shown to target ACE2 at the EC surface^28^.

Our analysis of the blood samples from COVID-19 patients showed the surprising result that the ACE2 found in the blood was a truncated form of the ectoenzyme. This observation was consistent with our result of the ACE2 extracted from the HUVECs. The level of this truncated form in the patient’s blood samples increased with the severity of the disease. Indeed, in severe cases of SARS-CoV-2 infection, there was one fragment of ACE2 that was not present in the blood samples from patients with mild disease. ACE2 functions as an enzyme in the renin-angiotensin system (RAS). The RAS plays a significant role within the physiology and pathophysiology of cardiovascular and renal systems. ACE2 has been proposed as an emerging biomarker of cardiac disease. Circulating levels of ACE2 may also have a prognostic role in monitoring COVID-19 infection, since it was found to be correlated with the severity of COVID-19 and a predictor for mortality^28^. Therefore, an increased level of ACE2 could result from increased ACE2 shedding due to the lysis of ACE2-expressing cells during due to severe lung infection^29^. Because the circulating ACE2 level is usually low in healthy donors, it can be suggested that elevated circulating ACE2 may predispose patients to severe disease^30^.

Furthermore, our results showed that the fully glycosylated extracellular domain of human VE-cadherin (sVE) was a substrate for ACE2. This is in agreement with our measurements of soluble forms of the ectodomain of VE-cadherin in the peripheral bloodstream of the COVID-19 patients. We have previously identified a soluble VE-cadherin exhibiting an apparent molecular weight of 90 kDa present in blood from patients with diseases such as rheumatoid arthritis, angioedema, and cancer^8,9,13^. In our reported results, for the first time we found three different forms of soluble VE-cadherin with apparent molecular weights of 70, 62, and 54 kDa in the blood of patients presenting severe COVID-19 disease. In blood samples from patients with mild COVID-19 disease, we detected only one form of soluble VE-cadherin with a molecular weight of 54 kDa. Of great significance to these observations is that ACE2 is a metallopeptidase orientated with its catalytic site facing the extracellular space as an ectoenzyme and therefore can cleave circulating peptides. We have identified that the extracellular domain of VE-cadherin contains these two amino acid sequences at the positions ^256^P-F^257^ and ^321^PMKP-^325^L for ACE2 substrate recognition. The putative site for ACE2 cleavage (^321^PMKPL^325^) was located in EC3 ectodomain of VE-cadherin, which could produce the 62 kDa form that we measured. The 70 kDa form could correspond to the site for Kallikrein found in patients with Hereditary angioedema. We cannot exclude the involvement of other proteases such as neutrophil elastase, which was recently published in COVID-19 patients^31^. Moreover, the protease elastase was already described to hydrolyze the ectodomain of VE-cadherin as well as serine proteases such as proteinase 3, cathepsins, and ADAMs from neutrophils^32^. Therefore, taking into account the local pulmonary environment and the damage to other organs in COVID-19, we hypothesize that the combined actions of ACE2 and/or other proteases can directly target VE-cadherin after viral infection, a process that might be at the origin of vascular lesions. Thus the analysis of the apparent molecular weights of sVE identified in the blood from COVID-19 patients could predict the protease involved in the cleavage process and consequently provide therapeutic medications to maintain EJ integrity.

## Conclusions

The recent clinical reports^3,4^ coupled with previous publications^4,5^ provide strong evidence to consider that the EJ structure is a central target that leads to EC damage, endothelialitis, coagulopathy and microcirculation dysfunction in SARS-CoV-2 infection. According to the Zurich team, COVID-19 causes vascular inflammation that can occur in different organs^3^. To the best of our knowledge, we present here the first proof of concept that identifies VE-cadherin as a target of ACE2. The consequent cleavage of the extracellular domain of VE-cadherin provides an explanation for degradation of the EJ and hence the vascular lesions observed in COVID-19 disease. Furthermore, the combination of in vitro and in vivo observations facilitated by measuring both ACE2 and soluble VE-cadherin (sVE) in blood, provides a mean to predict gradations in the severity of SARS-CoV-2 infection. The quantification of sVE might provide a blood test for personalized COVID-19 patient care, management and follow-up. This measurement of sVE provides an intriguing direction for the clinical management of COVID-19, and further investigations will be of major importance to determine the association between VE-cadherin, vascular endothelial dysfunctions and coagulopathies in long COVID-1933. Future fundamental research is needed to explore whether SARS-CoV-2 targets indirectly VE-cadherin through proteases or other activated pathways within the SARS-CoV-2 environment. Those different pathways would result in different sizes of the soluble extracellular fragments of VE-cadherin found in the blood, which can be used to inform the clinicians about the specific protease involved in the SARS-CoV-2 environment to damage the endothelium. This exciting possibility highlights the need for continuing efforts to elucidate whether the size of sVE fragments in the blood of COVID-19 patients might be of prime significance for clinicians to have quantifiable information on which to base decision-making to precisely target the actors of “infection” and “damage” caused by COVID-19 to allow “personalised” medicine.

## Authors’ Contributions

MB, CG, LBouv, and OG conducted the experiments. LBoui, DKM and IV participated in the design, interpretation of the studies and analysis of the data. All the authors reviewed the manuscript; LBoui, DKM and IV wrote the manuscript.

## Acknowledgments

The authors are grateful for the assistance of nurses of the Internal Medicine department for blood sampling.

## Ethical Approval

Studies involving human subjects have been approved by the local Ethics Committee (Grenoble University Hospital).

